# Molecular Insights into the Differential Dynamics of SARS-CoV-2 Variants of Concern (VOC)

**DOI:** 10.1101/2021.10.22.465272

**Authors:** Nabanita Mandal, Aditya K. Padhi, Soumya Lipsa Rath

**Affiliations:** National Institute of Technology, Warangal, Telangana, 506004, India; Laboratory for Structural Bioinformatics, Center for Biosystems Dynamics Research, RIKEN, 1-7-22 Suehiro, Tsurumi, Yokohama, Kanagawa 230-0045, Japan

**Keywords:** VOCs, Delta, SARS-CoV-2, Molecular Dynamics, RBD, ACE2

## Abstract

Severe Acute Respiratory Syndrome Coronavirus 2 (SARS-CoV-2) has affected the lives and livelihood of millions of individuals around the world. It has mutated several times after its first inception, with an estimated two mutations occurring every month. Although we have been successful in developing vaccines against the virus, emergence of variants has enabled it to escape therapy. Few of the generated variants are also reported to be more infectious than the wild-type (WT). In this study, we analyze the attributes of all RBD/ACE2 complexes for the reported VOCs, namely, Alpha, Beta, Gamma, and Delta through computer simulations. Results indicate differences in orientation and binding energies of the VOCs from the WT. Overall, it was observed that electrostatic interactions play a major role in the binding of the complexes. Detailed residue level energetics revealed that the most prominent changes in interaction energies were seen particularly at the mutated residues which were present at RBD/ACE2 interface. We found that the Delta variant is one of the most tightly bound variants of SARS-CoV-2 with dynamics similar to WT. High binding affinity of RBD towards ACE2 is indicative of an increase in the viral transmission and infectivity. The details presented in our study would prove extremely useful for the design and development of effective therapeutic strategies for the emerging variants of the virus.

## Introduction

Severe Acute Respiratory Syndrome Coronavirus 2 (SARS-CoV-2), is one of the largest known RNA viruses with a single-stranded RNA ranging between 26,000 to 32,000 bases.^1^ RNA viruses have a higher mutation rate compared to the DNA viruses, thereby, reflecting a higher replication fidelity of the DNA-dependent DNA polymerases over that of the RNA-dependent RNA polymerases.^2–4^ Additionally, positive-sense single stranded RNA viruses, such as SARS-CoV-2, have a much higher mutation rate than the negative-sense single stranded RNA viruses.^5^ The estimated rate of mutation reported is two per month for SARS-CoV-2.^6^ Mutation is also one of the primary generators of diversity among the genomes, including the viral genomes.^6,7^ Thus, we are observing an emergence of variants of SARS-CoV-2 since the first incidence of the Coronavirus disease 2019 (COVID-19).^8,9^

Particularly, mutation in the gene which encodes the Spike glycoprotein has been reported to be very high.^10^. In Spike protein, mutations can affect the transmission rate of the virus and also the disease outcome.^11^ The first reported mutation of the Spike protein was D614G mutation, which was found to enhance the SARS-CoV-2 transmission.^11^ The N501Y mutation increased the affinity of the Spike protein for its receptor, Angiotensin Converting Enzyme 2 (ACE2), thereby increasing the chances of viral transmission.^12^ Mutation E484K is known to contribute to the evasion of antibody neutralization.^13^ D796H and H655Y mutations that are present in the Spike protein, are associated with reduced affinity towards the neutralizing antibodies.^14^ The World Health Organization (WHO) has recently assigned different labels for the generated variants of SARS-CoV-2. They can be broadly separated into two categories, namely, the Variants of Concern (VOC) and the Variants of Interest (VOI).^15^ VOC has increased transmissibility and severity of (COVID-19) compared to VOI. There are four recognized VOCs (and several emerging variants while this work was being carried out); Alpha, Beta, Gamma and Delta. Alpha (strain B.1.1.7) is estimated to be 40– 80% more transmissible than the Wild-type SARS-CoV-2; Beta (B.1.351, B.1.351.2, B. 1.351.3) has three mutations in the receptor-binding domain in the Spike glycoprotein of the virus: N501Y, K417N, and E484K respectively where, two of them (E484K and N501Y) mutate at the receptor-binding motif (RBM); Gamma (P.1, P.1.1, P.1.2) has about ten mutations in the Spike protein, where three mutations namely N501Y, E484K and K417T are of particular concern occurring at the RBM; and Delta (B.1.617.2, AY.1, AY.2, AY.3) where mutations occur at RBD regions T478K, P681R and L452R. The Delta variant is of particular interest because it evades the neutralizing antibodies and also induces higher cell-cell fusion in the respiratory tract, contributing to the chance of higher pathogenicity.^16^

All the mutations observed in the VOC were primarily located in the RBD ranging from residue 333-527 of the Spike protein.^17^ The RBD of the SARS-CoV-2 Spike protein interacts with a human ACE2 receptor. This receptor is found on the lung alveolar epithelial cells and plays a primary role in protection against lung injury in humans.^18^ The RBD consists of a N-terminal and C-terminal domain and also a Receptor Binding Motif or RBM. The RBM is responsible for stable interaction with the host receptor, ACE2. This RBM consists of four loops which are divided by two small beta-strands.^19^ Several studies have shown that the difference in the sequence of RBD between SARS-CoV-1 and SARS-CoV-2 have increased the binding affinity of the SARS-CoV-2 RBD towards ACE2.^20^ It is therefore essential to learn how these mutations impact the association of the RBD with the ACE2.

Very recently, the cryo-EM structure of the Delta Variant has been solved,^21^ however, the differences in structures and binding are yet to be unraveled. Moreover, while this manuscript was being communicated, an experimental study on some of the emerging SARS-CoV-2 variants is reported,^22^ which corroborates our findings and offers an opportunity to critically assess our MD results. In the present study, we used molecular modelling tools to model the RBD domains of all the reported VOCs. Subsequently, we compared the generated models of the VOCs in the RBD/ACE2 complex with the Wild-type by using extensive molecular dynamics (MD) simulations and identified several key features which result in differential activity of the VOCs. This study provides mechanistic and molecular insights of the VOCs and would prove crucial for understanding the structure-function relationship as well as in the development of effective therapeutic strategies for the emerging variants of SARS-CoV-2.

## Materials and Methods

### Protein systems and setup

The crystal structure of the Spike protein RBD associated with ACE2 was taken from Protein Data Bank (PDB ID: 6LZG) as the starting structure.^23^ Although 13 residues were missing from the N-terminal of ACE2, the N-terminal residues do not directly interact with the RBD,^23^ hence they were not modelled. This structure was considered as the Wild-type system. The variants namely, Alpha (P.1), Gamma (B.1.1.7), Beta (B.1.351) and Delta (B.1.167.2) were generated by mutating specific residues in Wild-type Spike protein after aligning the RBD sequences shown in Figure 1. Modeller 10.1 molecular modelling suite was used to generate the new models based on the Wild-type RBD/ACE2 template.^24^ We also checked the initial structures of the generated variants by observing the distribution of their phi-psi angles and other stereochemical properties by using the PROCHECK^25^ suite of programs. To understand the effect of mutations on the structure and dynamics of the Spike-ACE2 complex, we performed all-atom MD simulations of Wild-type (WT) and four variants of Spike-RBD/ACE2 in triplicates.

**Figure 1.**
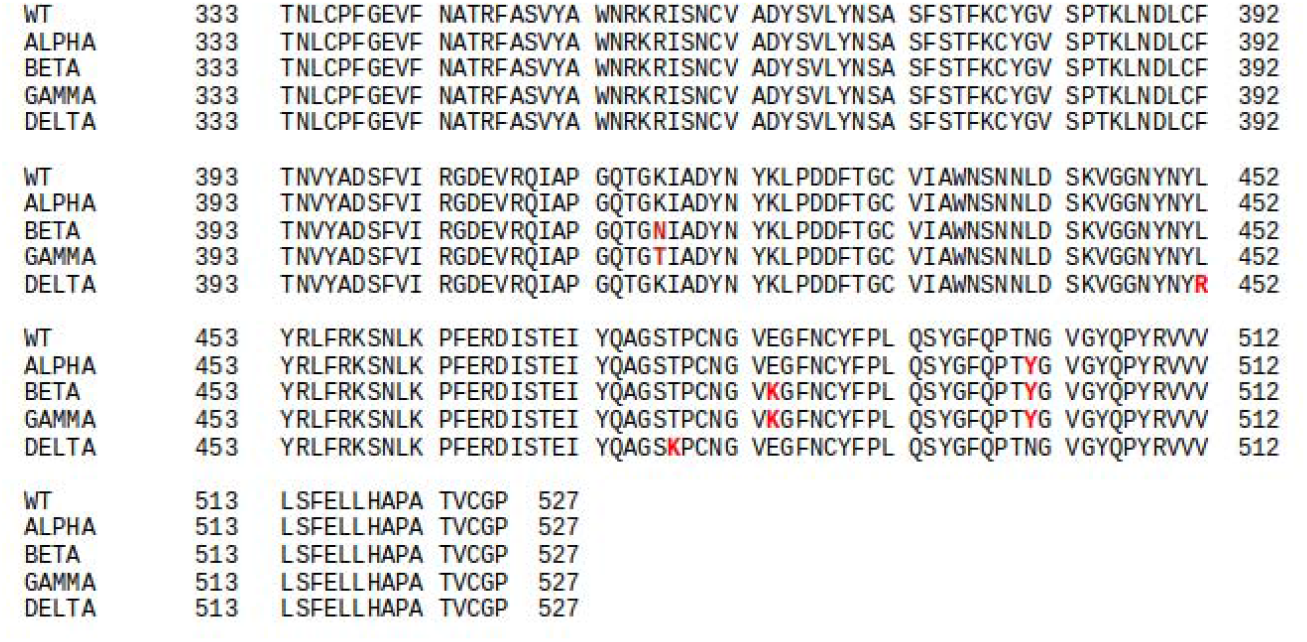
Complete sequence of wild-type (WT) and the Variants of concerns (VOCs). The mutated residues are highlighted in red.

### Molecular Dynamics Simulations

Atomistic MD Simulation was carried out using Gromacs MD Simulation package^26^ using CHARMM36 force field parameters.^27^ Each system was subjected to energy minimization in steepest descent and then in conjugate gradient for 2000 steps. After initial relaxation, a cubic simulation box consisting of three-site TIP3P water molecules and neutralizing ions was created for the systems.^28^ The box dimensions were 10 x 10 x 10 □. For charge neutralization 24 ions were randomly placed by replacing the corresponding solvent molecules. Subsequently, energy minimization and thermalization was performed to avoid any bad contacts which might have been created due to the mutations and addition of water and ions. Periodic boundary condition was implemented during simulation. The systems were gradually heated from 0 to 310 K for 200 ps. Then the systems were equilibrated at 310 K in NVT ensemble using modified Berendsen thermostat^28^ for about 500ps and then equilibrated in NPT ensemble using 1 atmospheric pressure using Parrinello-Rahman barostat^28^ for 1 ns. A time step of 2 fs was used for all the equilibration and subsequent production runs. After the convergence of potential energy and density, production simulation was carried out for the Wild-type and VOCs for 100 ns in NPT where the coordinates were saved at the interval of every 1000 ps. Particle-mesh Ewald method was used to treat the long-range electrostatic interactions. VMD and Pymol were used for visualization of the trajectories.^30,31^ All the analyses were carried out using Gromacs tools^26^.

### Binding energy calculation between RBD and ACE2

The binding energy between RBD and ACE2 for WT and VOCs (P.1, B.1.1.7, B.1.351 and B. 1.167.2) were computed by using the Molecular Mechanics/ Poisson Boltzmann Surface Area (MM/PBSA) employed in the g_mmpbsa tool of GROMACS.^32^ In this methodology, the binding energy of the target-ligand or protein-protein is typically defined as

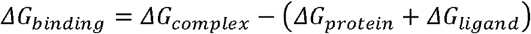

where ΔG_protein_, ΔG_complex_, and ΔG_ligand_ represent the total free energies of the complex, the ligand, and the protein also separately in the solvent, respectively. Further, the free energy of the separate entity is represented as

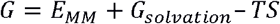

where E_MM_ stands for the average molecular mechanic’s potential energy in the vacuum, G_solvation_ denotes the free energy of solvation. TS stands for the entropic augmentation to the free energy in a vacuum, here S and T denote the entropy and temperature, respectively. The E_MM_ consists of nonbonded and bonded terms, including torsion, bond angle and electrostatic (E_elec_) and the Van der waal (E_vdw_) interactions, respectively & the solvation free energy, G_solvation_ takes both electrostatic and non-electrostatic (G_polar_ and G_nonpolar_) components. The binding free energy for the complexes was calculated from 50 snapshots over the last 10 ns of the simulation trajectories. All the systems were stable during this period.

### Contact analysis

To understand the intermolecular interactions formed between RBD and ACE2 for WT and all the VOCs, contacts (hydrogen bonds and salt bridges) were computed and analyzed from the last 10 ns MD simulated trajectories using GetContacts.^33^ The hydrogen bonds were shown in a clustergram to make the interpretation clear for visualization.

## Results and Discussion

### A. Modeling and simulation of Wild-type and VOC’s RBD/ACE2 complex

The initial coordinates of the RBD/ACE2 complex were taken from the crystal structure (PDB ID: 6LZG)^34^. As shown in Figure 1 and Figure S1, the mutations of the VOCs occur at specific sites on the RBD of the Spike protein. Due to the unavailability of structural information of all the variants during the beginning of our study, we used Modeler10.1 molecular modeling suite^35^ to generate four energetically stable structures RBD/ACE2. The stability of the structures was verified through the DOPE score of Modeller10.1 (data not shown). The Ramachandran plot of the generated models did not show drastic changes as expected from a static model (Figure S2). Notably, these mutations occur close to the RBD/ACE2 interface. It is therefore fascinating to investigate how Spike-ACE2 interaction and dynamics might be affected due to mutations.

To begin with, we ran atomistic MD simulations in triplicates for WT, Gamma, Beta, Alpha and Delta variants for 100 ns at normal temperature and pressure (NPT) conditions (Table 1). Figure S3 shows the root mean square deviation (RMSD) of the five systems over the simulation run time. From the figure it is evident that all the systems have similar RMSD values, however, Delta was found to be remarkably stable than the others. All the systems had reached stability around 50ns of the simulation with the time-average RMSDs lying within the range of 0.25-0.3 nm. The average RMSD calculated from the three replicates did not show much difference in values further substantiating our observations (Table S1)

**Table 1.** Variants under study and their abbreviation used the article

| Serial No. | Variant | Abbreviation |
| --- | --- | --- |
| 1 | Wild-type | WT |
| 2 | B.1.1.7 | Alpha |
| 3 | B.1.351 | Beta |
| 4 | P.1 | Gamma |
| 5 | B.1.167.2 | Delta |

Once the systems reached stability, we further analyzed the fluctuations of the C□ atoms of the RBD and ACE2 proteins separately. Figure 2a shows the Root Mean Square Fluctuation (RMSF) of ACE2 protein. The ACE2 protein mainly plays a role in the cardiovascular system and in lungs, however its presence is also seen in other organs.^36^ Further reviews on its role in various metabolic pathways are described elsewhere.^37^ Structurally, ACE2 is an alpha helical protein,^37^where the N-terminal helical part of the protein is the primary site of interaction with the RBD of the Spike protein of SARS-CoV-2.^38^ The overall binding mode of ACE2 with both SARS-CoV-1 and SARS-CoV-2 is known to be similar.^39,40^ In comparison with the WT, we can clearly observe increased fluctuations in the Alpha, Beta and Gamma complex (Figure 2 and S4). However, ACE2 receptor in Delta show fluctuations similar to that of the WT protein. The N-terminal domain and residues Glu329, Asn330 and Lys353 are mainly involved in building H-bonded/salt bridge interactions between the proteins in RBD/ACE2 complex, hence increased fluctuations would indicate a weaker complex formation (Figure 2a). We also compared the dynamics of the RBD as well as RBM in all the systems, where slightly higher peaks were observed for Alpha, Beta and Gamma but the Delta system was remarkably stable (Figure 2b). It is generally expected that mutations at residue level would alter the protein dynamics, however, even though mutations existed in Delta, we could not find drastic changes in the fluctuations of the CA atoms.

**Figure 2.**
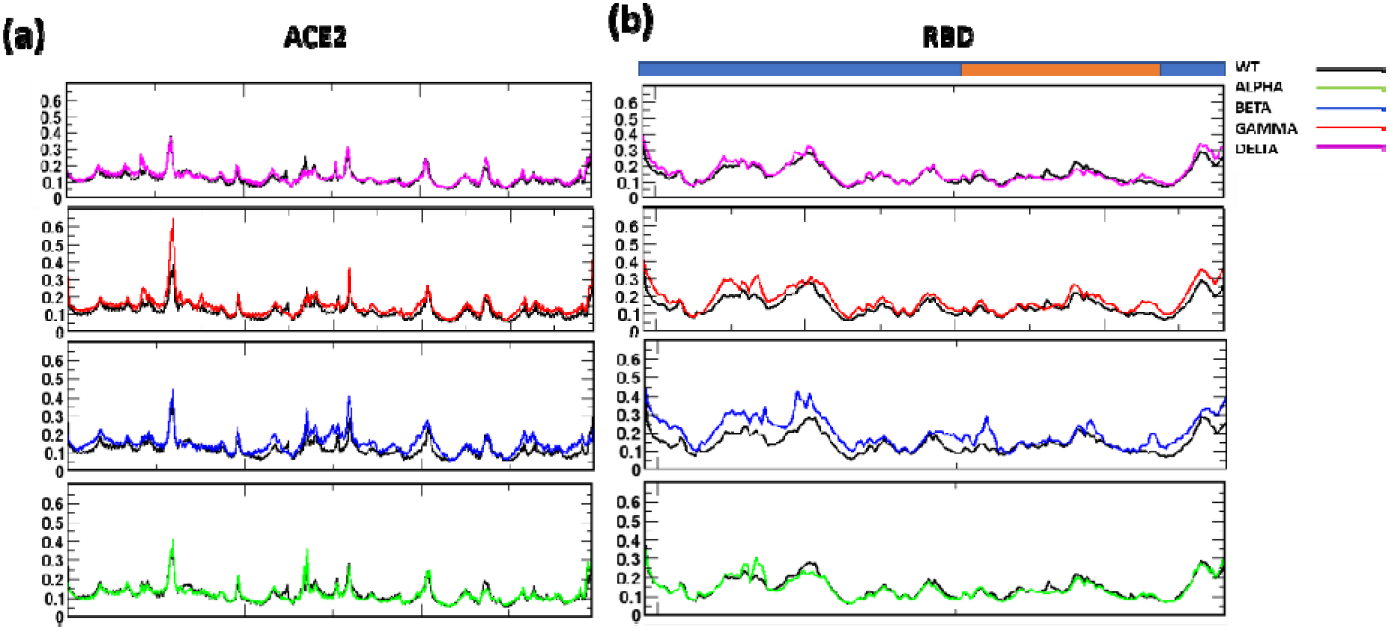
RMSF of (a) ACE2 and (b) RBD with respect to WT complex. In both ACE2 and RBD, the residues from B.1.1.7 and B.1.617.2 show more stability as well as similarity with the WT complex. The RBD region is shown as cyan bar in (b), where the RBM is highlighted in orange.

### B. Difference in structure and dynamics of the VOC’s

Figure 3 shows the time-averaged structures of the variants superimposed on the WT ACE2 receptor, to verify the relative orientation of the RBD with respect to ACE2. The interface formed by the N-terminal helices of ACE2 and the RBD were compared among the systems. From the figure we observed that the interface loop region of RBD and helices of ACE2, in the Gamma complex, has moved relatively away when compared to WT (Figure 3c). On the other hand, Alpha superposes very well with the WT (Figure 3a). Beta as well as Delta, however, appear to have moved closer to the ACE2 complex (Figure 3b, d). Subsequently, we analyzed the overall RBD/ACE2 complex after superimposition. Here, we found that although the orientation of RBD in Alpha and Gamma complexes were similar (Figure 3e, g), Beta as well as Delta shows a stark difference in the orientation of RBD (Figure 3f, h). In both the complexes we observed a relative rotation of the RBD w.r.t. ACE2. This significant shift in the protein orientation would influence its dynamics as well as residue positions and interactions in the protein-protein complex. To elucidate the change in the dynamics of the protein, we used Principal Component Analysis (PCA) on the generated trajectories and studied the dynamics of the RBD with respect to the ACE2 receptor. PCA captures the dominant motions of the protein by using a set of eigenvectors. The most significant motion of a protein can be captured with the eigenvector with the maximum eigenvalue. We calculated the distribution of PC1 and PC2 for all the five complexes (Figure S5) and observed a relatively wide distribution of the Beta and Gamma RBD/ACE2 complexes when compared to the WT, Alpha or Delta indicating an increase in complex dynamics.

**Figure 3.**
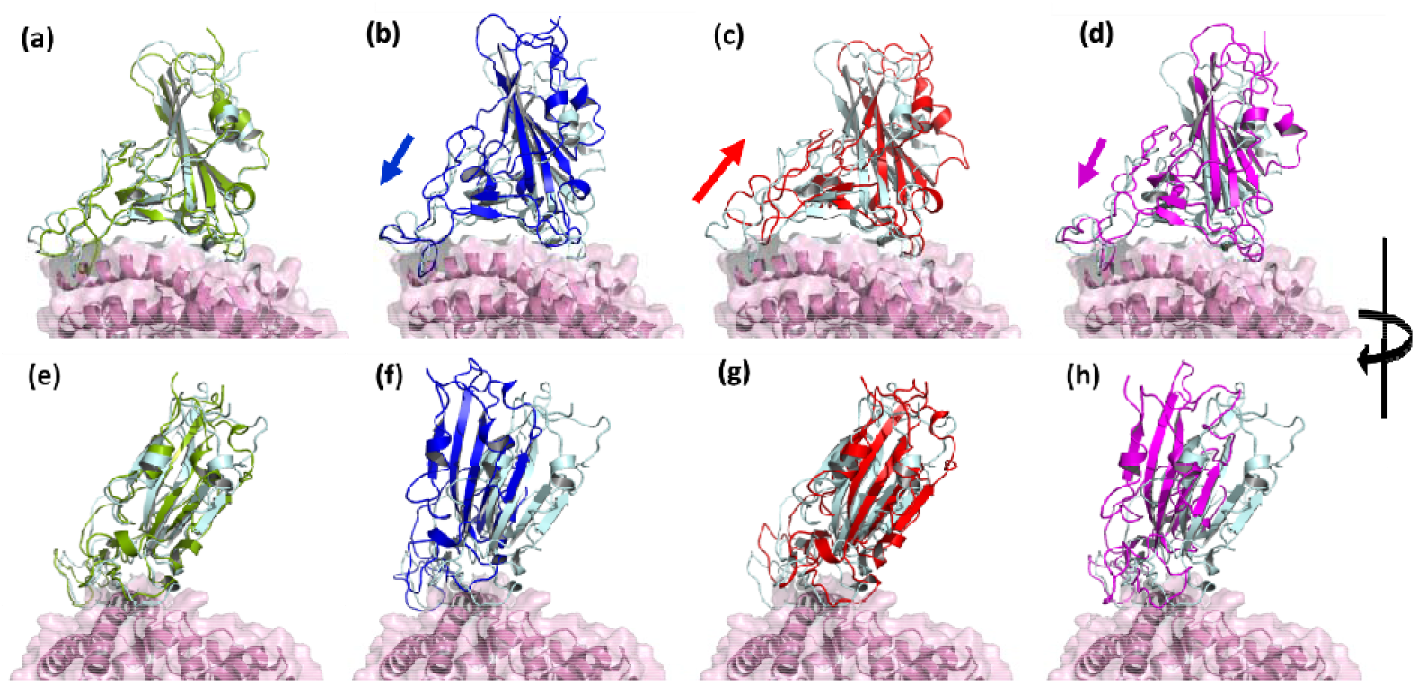
Superimposed structure of WT and VOCs showing the relative variation in the receptor binding domains. Interfacial region of RBD and ACE2 in (a) Alpha, (b) Beta, (c) Gamma and (d) Delta. When the structure is rotated 90° in (e) Alpha, (f) Beta, (g) Gamma and (h) Delta. Relative displacement was observed in Beta and Delta. (WT: RBD in cyan, ACE2 in pink; Alpha RBD in green; Beta RBD in Blue; Gamma RBD in Red; Delta RBD in Magenta).

Subsequently, we calculated the principal components on the backbone C□ atoms of RBD and ACE2 separately.^41^ The Gibbs Free energy landscape was then constructed as a function of the PC1 and PC2 coordinates. The highly stable protein conformation is shown in red, other low energy states are colored either in blue, green, or cyan. In the WT complex, the ACE2 was confined to a single cluster whereas, RBD explored two separate clusters (Figure 4a). This indicates that RBD can exist in two different conformations after being bound to ACE2. Although not much changes could be seen in the binding of RBD, the difference in clusters can be mainly attributed to the loop dynamics. When we compared the variants, we found that a greater number of conformational states of the ACE2 in variants Alpha and Beta (Figure 4 b, c), however, the ACE2 receptor of Gamma and Delta explored the same low energy conformations (Figure 4 d, e). This is expected since the mutations have primarily taken place in the RBD, which is devoid of large portions of the Spike protein. Moreover, despite the amino acid substitutions, the dynamics of the ACE2 receptor doesn’t get influenced significantly. Later, we observed the dynamics of the RBD domain, which is the prime site of variation. Although the RBD of all the systems explored two different clusters, Alpha and more prominently Gamma show remarkable differences in the protein dynamics (Figure 4b, d). While, Alpha shows a single cluster with the low energy state of the protein, in Gamma the two different clusters are relatively shallow with more scattered low energy states. Surprisingly, despite observing significant changes in the superposed structures (Figure 3), the dynamics of Beta and Delta of both RBD and ACE2 were similar to the WT complex (Figure 4a, c, e). Thus, overall, our results reveal stark similarity in the dynamics of the RBD/ACE2 complex between WT, Beta and Delta, but Alpha and Gamma show differences in the energetically stable states. We also checked for the trace values from the covariance matrix that was generated from the PCA. The trace values are correlated with the total variance in the values of eigenvectors where higher values indicate more variation.^42^ For ACE2 in WT, Alpha, Gamma, Beta and Delta the values were found to be 6.34 nm^2^, 6.61 nm^2^, 6.08 nm^2^, 6.09 nm^2^ and 5.98 nm^2^ respectively. Similarly, for RBD it was found to be 1.02nm^2^, 1.16 nm^2^, 1 nm^2^, 1.08 nm^2^ and 0.97 nm^2^ respectively. Thus overall, the maximum flexibility was observed for the Alpha complex in both ACE2 and RBD and the Delta had the least flexibility among all the systems under study. This indicates that the Delta was rather a tightly bound complex when compared to the other systems including the WT. This instigated us to check the total binding energies of RBD/ACE2 in all the systems.

**Figure 4.**
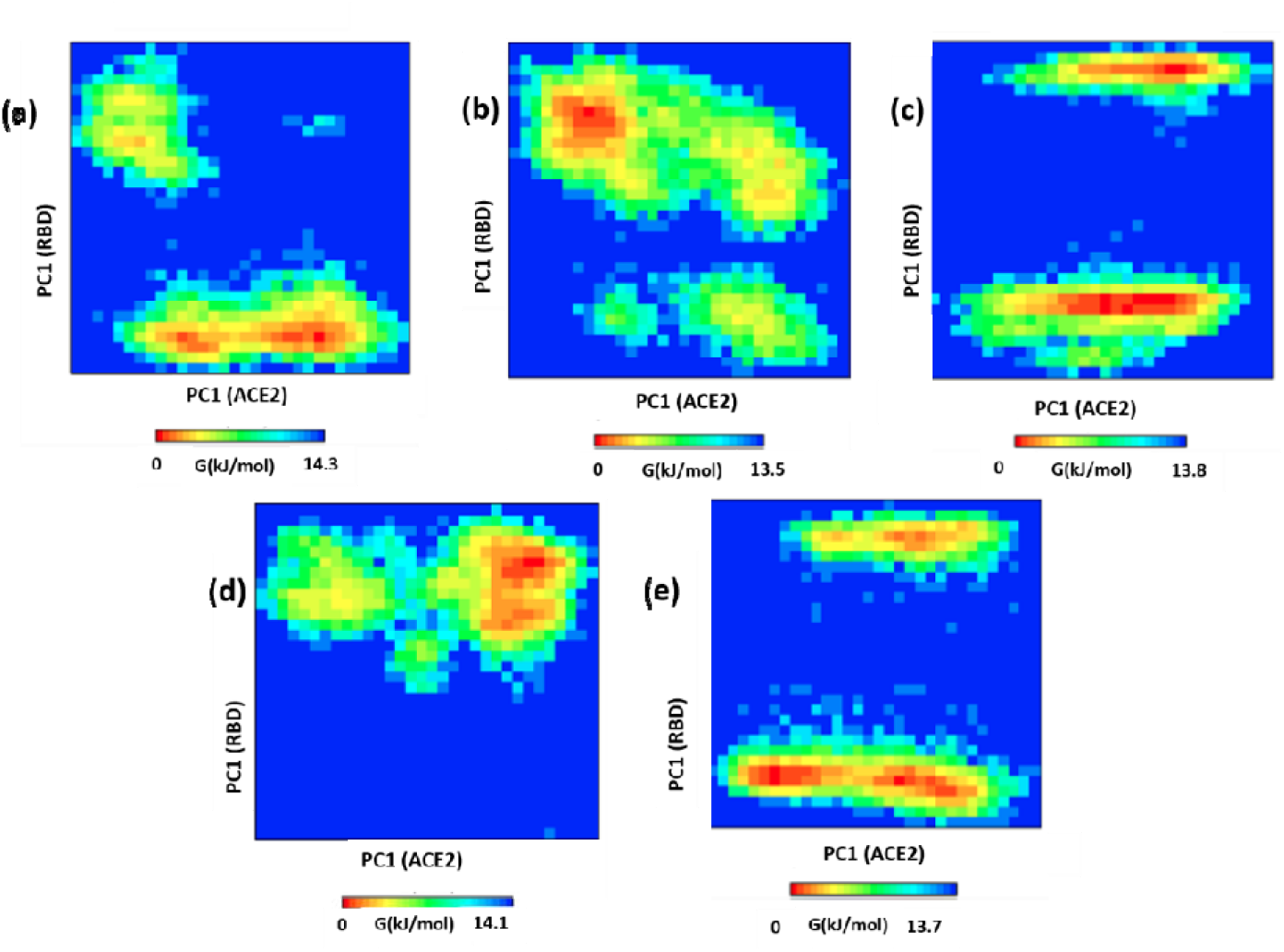
Free energy landscape of the WT and VOCs. The Gibbs free energy landscape was constituted from the simulated structures on the plane defined by the first principal component of ACE2 and RBD in (a) WT, (b) Alpha, (c) Beta, (d) Gamma and (e)Delta. The state with the lowest energy is coloured in red. The two clusters formed by RBD are shown in green and magenta cartoons. The clusters are formed primarily due to loop dynamics.

To explore the rationale behind the differential dynamics, we used the MM/PBSA to calculate the total binding energies of the protein complexes (Table 2). MM/PBSA is a widely used technique used for binding free energy calculations between protein-protein or protein-ligand systems. We used the last 10 ns of simulated trajectories of all the systems for MM/PBSA based binding energy calculations, where the systems were mostly stable. Table 2 shows the trend of the binding energies of the five systems under study. Accordingly, it is seen that the binding of Delta>Gamma>Beta>WT>Alpha. Upon comparison of the binding energies, we find Delta to have the highest interaction energy and Alpha has the least. We also see large changes in the electrostatic interactions among the complexes. Although there is an increase in electrostatic interaction energy which mainly accounts for the rise in the total binding energy of Gamma, an increase in the polar solvation energy indicates higher solvent interaction of the protein, which is in accordance with the changes observed in Figure 3a. Similarly, Alpha shows a significant increase in the solvent accessible surface area (SASA) energy, which hints towards the difference of the accessibility of the protein to the solvent. An increase in the SASA value here indicates conformational change in the RBD/ACE2 complex. From Table 2, it is evident that Beta and Delta are more compact when compared to other complexes which are similar to the trace values observed in PCA. Thus, it was seen that the binding energies of Delta is the highest and Alpha the least among all the five systems under study.

**Table 2.** Interaction energies between RBD and ACE2 in WT and VOCs

| List of VOCs | Van der Waal energy (kJ/mol) | Electrostatic energy (kJ/mol) | Polar solvation energy (kJ/mol) | SASA energy (kJ/mol) | Total Binding energy (kJ/mol) |
| --- | --- | --- | --- | --- | --- |
| WT | -363.775 ± 20.446 | -1232.614 ± 64.543 | 617.810± 161.796 | -43.889 ± 3.469 | -1022.467 ± 150.703 |
| Alpha | -334.114 ± 19.106 | -1010.046 ± 69.032 | 509.951 ± 82.835 | -74.747 ± 3.721 | -908.956 ± 91.026 |
| Beta | -282.277 ± 20.433 | -1659.780 ± 108.968 | 539.401 ± 156.941 | -36.157 ± 3.835 | -1438.813 ± 140.383 |
| Gamma | -344.279 ± 22.058 | -1628.128 ± 85.641 | 396.771 ± 168.090 | -42.888 ± 4.565 | -1618.525 ± 139.268 |
| Delta | -320.790 ± 20.134 | -1985.424± 67.699 | 558.329 ± 109.035 | -40.544 ± 3.624 | -1788.429 ± 99.618 |

### C. Interfacial residues significantly influence the stability of the RBD/ACE2 complex

Interfacial residues of proteins play a significant role in the association as well as in governing stability of the protein complexes. In the earlier studies by Spinello et al.,^43^ they compared the interface of SARS-CoV-1 with SARS-CoV-2 and found substantial differences in the interaction energies between residues of ACE2 and RBD. Here, we further calculated residue-wise contribution towards the total binding energy of the complex for both ACE2 and RBD by using MM/PBSA. Several of the residues were found to show drastic differences in their binding energies. We compared the energy of those residues that contributed >10 kJ/mol towards the binding energy of the complex. The interaction energies of RBD show radical changes in values for the Alpha complex. Here, almost all of the residues, except Glu484, contribute negligibly towards protein binding (Figure 5, Table 3a). We further noticed that Glu484 mutates into Lys484 in both Alpha and Beta which increases the binding energy to - 226.64+/−2.8 and −258.40+/−4.71 kJ/mol respectively. However, Glu484 in the WT and Gamma show highly repulsive energy values (212.50+/−1.1 and 199.02+/−0.84 kJ/mol respectively) indicating unfavorable interaction (Figure 5b). Similarly, substitutions L452R and T478K in Delta significantly increase the interaction energies by −199.57+/− 0.57 kJ/mol and −186.29 kJ/mol. Overall, it was found that five residues in Delta and Beta, four in WT and Gamma contribute the maximum in the binding of RBD to ACE2. Thus, overall mutations in Beta and Alpha comprises both unfavorable and favorable interactions. The changes observed for the Gamma variant w.r.t WT was not very significant. However, the mutations at L452R and T478K of the Delta tremendously contributed towards the interaction energy (around −386 kJ/mol) of RBD/ACE2.

**Figure 5.**
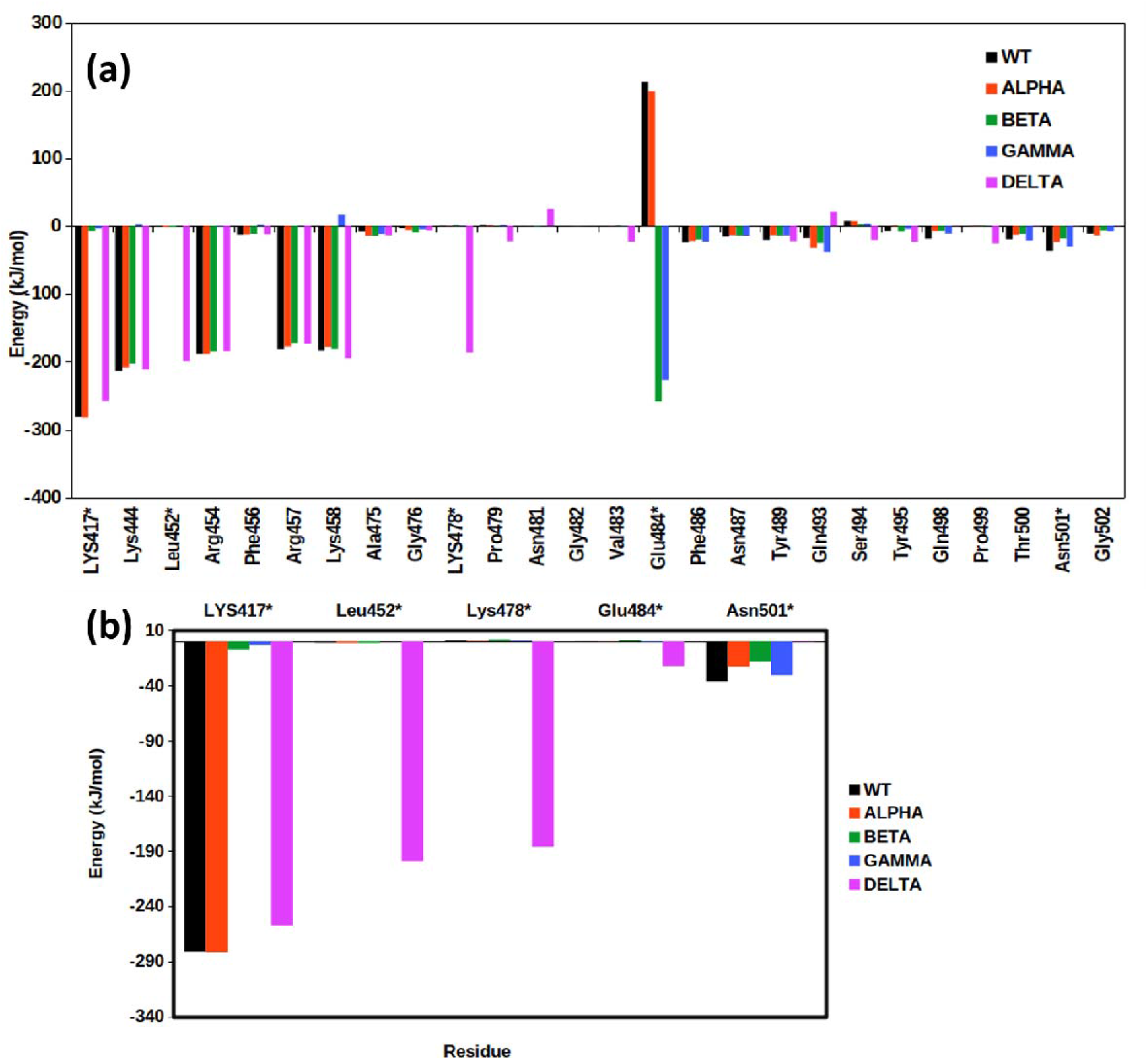
Residue level contribution towards the interaction energy. Contribution of (a) RBD and (b) mutated residues in RBD for all the five systems under study. The mutated residues are highlighted by a *.

**Table 3a.** Residue-wise interaction energies of the interfacial residues of RBD in RBD/ACE2 complex for the systems under study

| Residue | WT | Alpha | Beta | Gamma | Delta |
| --- | --- | --- | --- | --- | --- |
| LYS417* | -281.10 | -281.92 | -7.28 | -3.33 | -257.77 |
| Lys444 | -213.86 | -208.76 | -202.51 | 2.91 | -210.82 |
| Leu452* | -1.24 | -1.46 | -1.47 | -0.01 | -199.57 |
| Arg454 | -188.10 | -188.17 | -184.91 | -0.86 | -184.60 |
| Phe456 | -12.33 | -12.09 | -11.14 | 2.39 | -11.87 |
| Arg457 | -180.94 | -177.45 | -172.28 | 0.55 | -173.44 |
| Lys458 | -183.71 | -178.02 | -180.58 | 16.51 | -194.78 |
| Ala475 | -8.25 | -13.97 | -14.14 | -11.37 | -13.54 |
| Gly476 | -2.95 | -5.42 | -9.23 | -4.46 | -5.85 |
| Lys478* | 0.62 | 0.44 | 1.97 | 0.62 | -186.29 |
| Pro479 | 2.17 | 1.96 | 0.48 | 2.05 | -22.12 |
| Asn481 | -0.40 | 0.07 | -0.73 | -0.19 | 24.70 |
| Gly482 | -0.016 | -0.10 | -0.23 | 0.14 | 21.26 |
| Val483 | -0.43 | -0.21 | 1.08 | -0.48 | -22.49 |
| Glu484* | 212.50 | 199.02 | -258.40 | -226.64 | 0.11 |
| Phe486 | -23.92 | -21.67 | -19.56 | -22.89 | 0.02 |
| Asn487 | -15.48 | -13.38 | -13.91 | -14.37 | 0.01 |
| Tyr489 | -20.19 | -13.38 | -13.92 | -13.80 | -22.02 |
| Gln493 | -17.38 | -32.23 | -24.90 | -37.76 | 21.20 |
| Ser494 | 7.5175 | 7.3072 | 3.1298 | 3.8562 | -20.2757 |
| Tyr495 | -7.1823 | 1.012 | -8.1582 | -3.9698 | -23.1634 |
| Gln498 | -18.3783 | -7.2192 | -6.7676 | -11.0041 | -0.1788 |
| Pro499 | -0.4593 | 1.1247 | 0.8026 | -0.2207 | -25.5012 |
| Thr500 | -19.2305 | -12.4466 | -11.2612 | -20.8474 | -0.3149 |
| Asn501 * | -36.1766 | -23.2554 | -18.215 | -30.6474 | -0.2816 |
| Gly502 | -10.986 | -13.6278 | -5.9123 | -7.8995 | -0.1946 |

Subsequently, we checked if complementary changes take place in the ACE2 receptor of the complexes. Upon comparing the interaction energy of the ACE2 protein residues with RBD, calculated from the last 10ns of the trajectory (Table 3b), we found that in the WT complex except for residues Asp38, Tyr41, Tyr83, K31, Asp30 and Thr27, nearly all other residues show lower binding energies as compared to the VOCs. This indicates that mutations in the RBD impact the binding efficiency of ACE2 protein. Again, it was found that the residues of the Gamma variant show lower binding energies when compared to other variants. In Alpha, majority of energy was found to be contributed by charged-hydrophilic Glutamate and Aspartate residues, i.e., Glu22, Glu23, Asp30, Glu35, Asp38, Glu56, Glu57, Asp67 and Glu75. However, the complementary binding was absent in the RBD which reduces the overall binding efficiency. Both Beta and Delta ACE2 interfacial residues show significantly high energies of interaction, particularly around residues Glu22, Glu23, K31, Glu35, Glu37, Asp38, Glu56, Glu57, Asp67 and Glu75 in Beta and Glu22, Glu23, Asp30, Glu35, Glu37, Asp38, Glu56, Glu57, Asp67 and Glu75 in Delta (Figure S6).

**Table 3b.** Residue-wise interaction energies of the interfacial residues of ACE2 in RBD/ACE2 complex for the systems under study

| Residue | WT | Alpha | Beta | Gamma | Delta |
| --- | --- | --- | --- | --- | --- |
| Glu22 | -19.83 | -20.93 | -37.96 | -42.19 | -57.74 |
| Glu23 | -20.82 | -29.77 | -53.33 | -76.61 | -84.16 |
| Glu24 | -15.59 | -14.19 | -16.17 | -15.88 | -15.73 |
| Thr27 | -13.36 | -12.33 | -13.08 | -12.29 | -15.02 |
| Asp30 | -51.24 | -55.64 | -12.26 | -64.51 | -97.39 |
| Lys31 | -31.88 | -30.84 | -87.24 | 54.25 | 27.88 |
| His34 | -32.51 | -32.69 | -12.26 | -15.59 | -18.84 |
| Glu35 | -16.66 | -22.56 | -87.24 | -94.12 | -78.44 |
| Glu37 | -12.06 | -79.44 | -92.53 | -70.72 | -98.38 |
| Asp38 | -12.16 | -50.41 | -96.32 | -86.23 | -115.92 |
| Tyr41 | -12.06 | -12.83 | -12.65 | -7.63 | -10.51 |
| Gln42 | -12.16 | -4.78 | -7.16 | -2.77 | -8.45 |
| Glu56 | -24.91 | -17.40 | -37.09 | -33.98 | -40.61 |
| Glu57 | -35.16 | -24.04 | -48.73 | -45.94 | -54.36 |
| Asp67 | -21.14 | -16.88 | -53.30 | -47.74 | -48.85 |
| Glu75 | -6.01 | -8.051 | -104.24 | -74.85 | -49.54 |
| Tyr83 | -17.99 | -12.30 | -13.36 | -12.49 | -13.81 |

We superimposed the time averaged structure of the variants on WT complex to understand the relative loss and gain of interactions as observed from the contacts and MM/PBSA analyses. From Figure 6 we can clearly see that the residues from the RBD have moved farther away from ACE2 in the Alpha complex (Figure 6a), at a similar position in Gamma (Figure 6b) and closer in both Beta (Figure 6c) and Delta. (Figure 6d) ACE2 residues (Figure S7) on the other hand lie more or less at a similar position.

**Figure 6.**
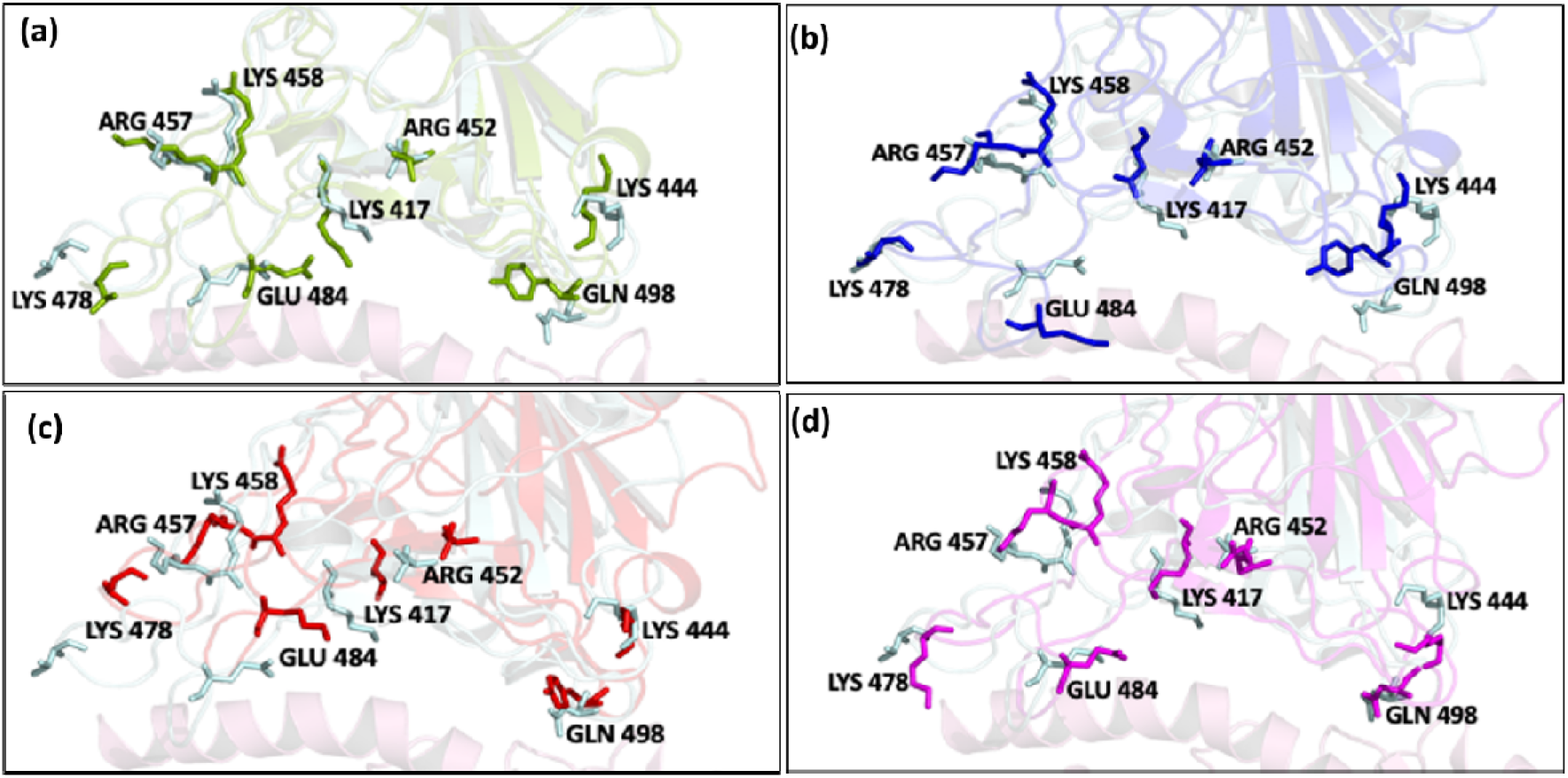
RBD interfacial residues primarily participate in interaction with ACE2. Superimposed images of WT with (a) Alpha, (b) Beta (c) Gamma and (d) Delta. Color scheme: WT (cyan), Alpha (green), Beta (blue) Gamma (red), and Delta (magenta). Negative displacement was prominently seen in Gamma while Beta and Delta moved towards the ACE2 receptor.

We proceeded to further validate our findings by constructing a map summarizing the probability of formation of salt bridges and hydrogen bonds among the five systems. For the analysis we considered the last 10ns of the stabilized trajectory. In two out of three complexes (i.e., Alpha and Beta) we didn’t observe any salt bridges. However, the salt bridges between Asp30 of ACE2 with Lys417 of RBD were found to be consistently present in WT, Delta and Gamma systems. This loss of interaction can be attributed to the mutation that occurs at residue Lys417 in both Alpha and Beta. We also checked for the difference in the H-bond interactions among the residues calculated over the last 10ns of the trajectory. Results show that most of the H-bond interactions reported in earlier studies^44,45^ were conserved in WT. ACE2 residue Tyr83 interacts with Asn487 of RBD across all the systems. There was a loss of H-bond between Asp30 of ACE2 with Lys417 of RBD in Alpha and Beta (Figure 7). We found Alpha complex to have the least number of H-bond contacts and WT and Delta to have the most H-bonded interactions with 7 and 5 H-bonded interactions for more than 50% of the time respectively (Figure 7). The two H-bonds lost in Delta were located at RBD residues Tyr505 and Thr500 with ACE2 residues Asp37 and Glu355 respectively. This change takes place due to the major conformational shift in RBD binding to ACE2 in the Delta complex. Gamma and Beta each have 4 H-bonds whose occupancy was more than 50% of the time. Very recently, the cryo-EM structure of the ACE2 bound RBD of the Delta variant was deposited in the Protein Data Bank. We made a comparison of the simulated structures with the Cryo-EM structure of the Delta variant (B.1.162.7) having the PDB ID:7V8B. ^46^The relative RMSD value was found to be only 1.7 □ which indicates similarity between both the structures (Figure S8). Further, upon comparison with a recent report on some of the emerging variants we found that substitutions in two of the residues i.e., Lys417 and Glu484K, show similar results. For instance, the salt-bridge breaking between Lys417 and Asp30 was seen in Alpha (Gamma) variant in both the systems. Similarly, we also observed enhanced binding energy upon E484K mutation in our studies. Additionally, we have also analyzed the interactions in the Delta (Delta) variant RBD with ACE2 where we have found it to be the most strongly binding variant of SARS-CoV-2 in the total binding energy calculation and observed tremendous contribution of mutated residues L452R and T478K towards protein-protein interaction. Thus, we observed that the interfacial residues, especially the mutated residues significantly contributed towards the stability of the RBD/ACE2 complex. In Delta variant this increase in the interaction energy leads to the formation of a compact RBD/ACE2 complex compared to the WT. This strong proteinprotein interaction along with dynamics close to the WT complex makes Delta one of the most tightly bound variants of SARS-CoV-2.

**Figure 7.**
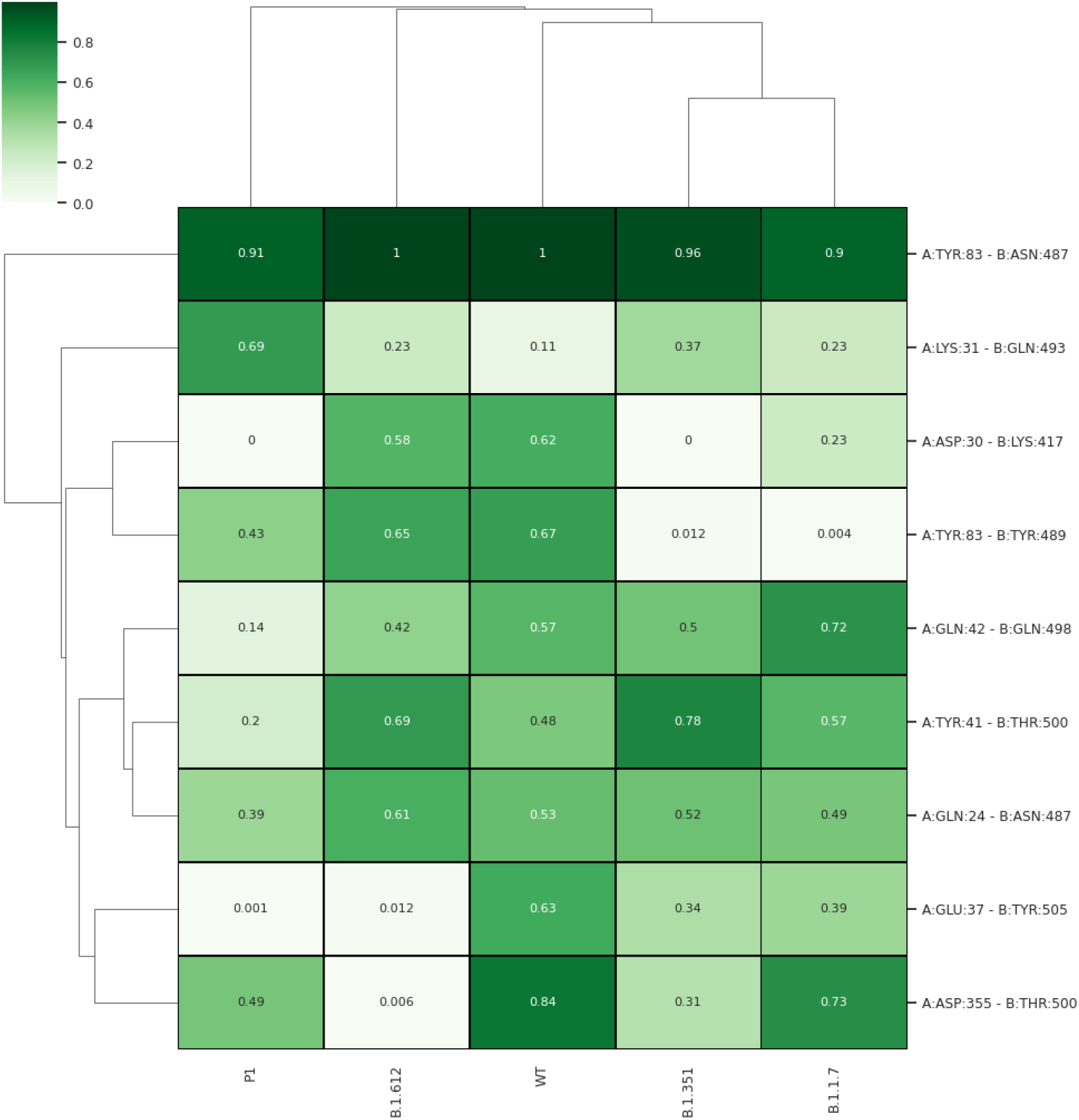
Computed hydrogen bonds between RBD and ACE2 for WT and VOC complexes. The gradient of green shows the frequency of interactions in the last 10 ns MD simulation trajectories. The contacts were shown as clustergram to make the interpretation clear for visualization.

## Discussion and Conclusion

COVID-19 pandemic caused by SARS-CoV-2 has affected millions worldwide. Although vaccines and antibodies have been developed against COVID-19, there is a constant effort to identify and tackle novel and emerging varieties of the virus. In such a condition, it becomes extremely important for us to understand the molecular level details of interaction of the variant with the host receptor. In the present study, we have used molecular modelling tools to model the RBD domains of the Spike protein of SARS-CoV-2 bound to the human ACE2 receptor. Subsequently, we compared the generated models in the RBD/ACE2 complex with the WT system by atomistic simulations. It was seen that at the interface region Gamma RBD was loosely bound to ACE2 when compared to Beta and Delta. Surprisingly, conformational changes could be observed in the binding mode of RBD in Beta and Delta after simulation, where we could observe relative rotation of RBD w.r.t ACE2. Protein dynamics of RBD and ACE2 show that Beta and Delta fluctuations correspond well with WT, unlike Gamma and Alpha. The Delta complex was also found to be the most compact system indicative of tighter complex binding. MM/PBSA analysis indicated drastic gain of interaction energy particularly in the Beta, Gamma and Delta. Overall, it was observed that electrostatic interactions play a major role in the binding of the complexes. Detailed residue level energetics revealed that the most prominent changes in interaction energies were seen particularly at the mutated residues. The mutations in Beta and Gamma were both a mix of unfavorable and favorable interaction but highly favorable for Delta variant. This increase in the interaction energy hints would lead to stronger association of RBD to ACE2 compared to the WT. The strong interaction energy coupled with dynamics similar to the WT complex makes Delta one of the most tightly bound variants of SARS-CoV-2. Comparison of the recently solved cryo-EM structure of the Delta (7V8B) revealed high structural similarities with the final time-averaged structures obtained after simulation. The high affinity of RBD and ACE2 is indicative of an increase in the viral pathogenicity. Therefore, the present study would prove extremely crucial for design and development of effective therapeutic strategies for the emerging variants of the virus.

## Acknowledgements

NM thanks MHRD Government of India for her Doctoral Scholarship. SLR thanks NIT Warangal for research seed grant (P1131) and National Energy Research Scientific Computing Center of the Ernest Orlando Lawrence Berkeley National Laboratory, a DOE Office of Science User Facility supported by the Office of Science of the U.S. Department of Energy under Contract No. DEAC02-05CH11231 and the Extreme Science and Engineering Discovery Environment (XSEDE). We are grateful to the COVID-19 HPC Consortium for providing resources and helping researchers work for a noble cause.

## Author Contribution

NM and SLR designed research; NM performed research; NM and AKP analyzed data; NM, SLR and AKP wrote the paper.

## Additional Information

### Supporting Information Available

Includes Figures S1-S8 and Table 1. This information is available free of charge *via* the internet

