## Supplementary Information for "Molecular Insights into the Differential Dynamics of SARS-CoV-2 Variants of Concern (VOC)"

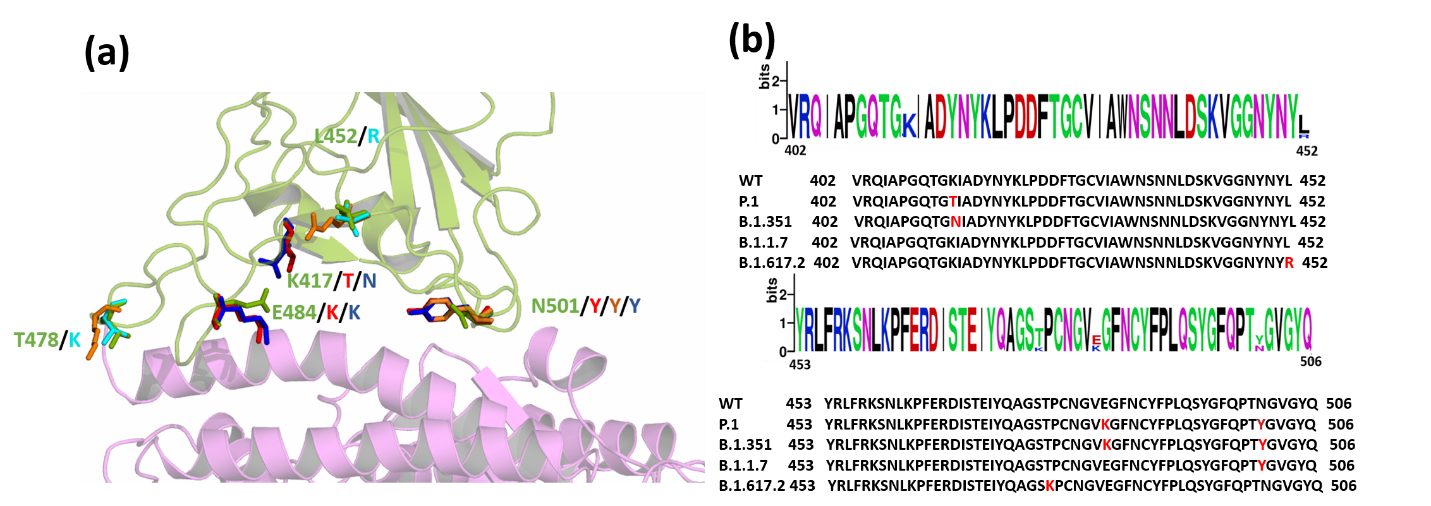

**Figure S1.** (a) Superimposed structure of the WT and the Variants highlighting the mutations in RBD. Wild type ACE2 receptor is shown in magenta and RBD in green cartoons. The mutations are shown as sticks. (WT: green, Gamma: red; Alpha: Orange; Beta: blue; Delta: Cyan) (b) Sequence alignment of WT and the VOCs. Mutated residues are highlighted in red in the sequence

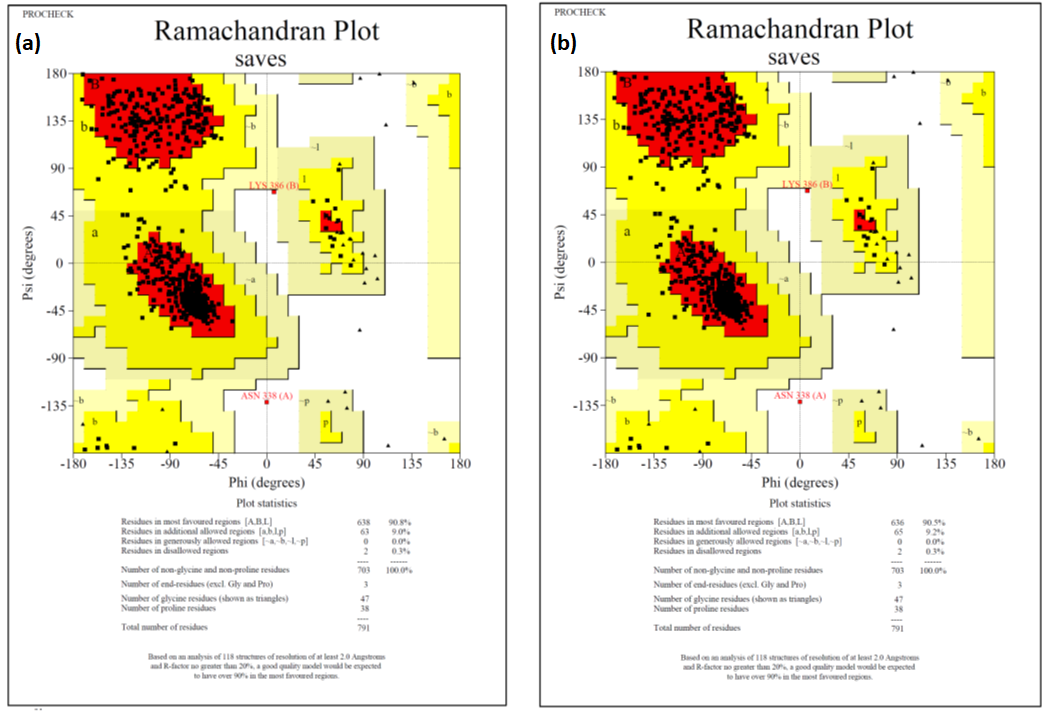

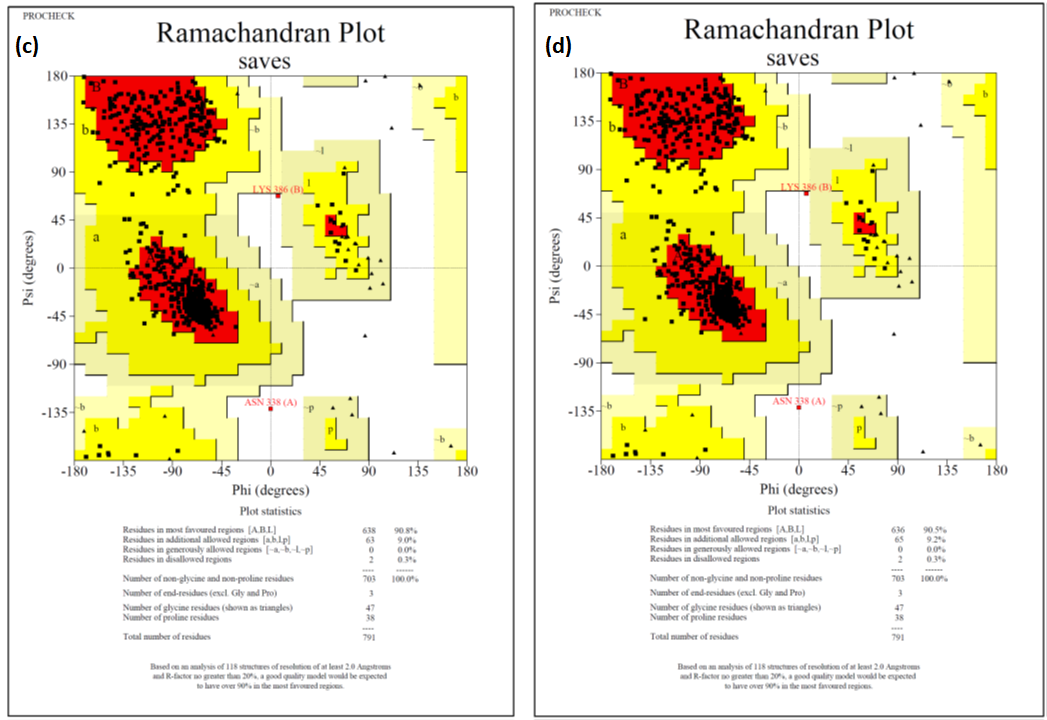

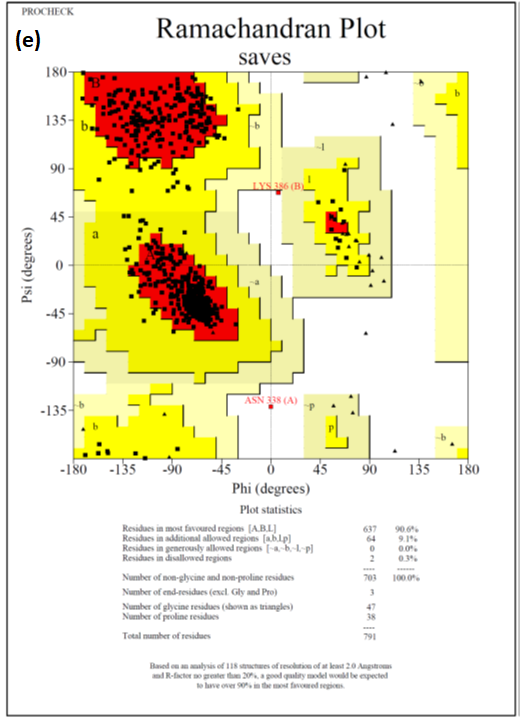

### Figure S2. Ramachandran plot generated by PROCHECK analysis displaying the stereochemical properties of WT and the four models that were generated showing structural similarity. (a) WT (b) Gamma (c) Alpha (d) Beta and (e) Delta.

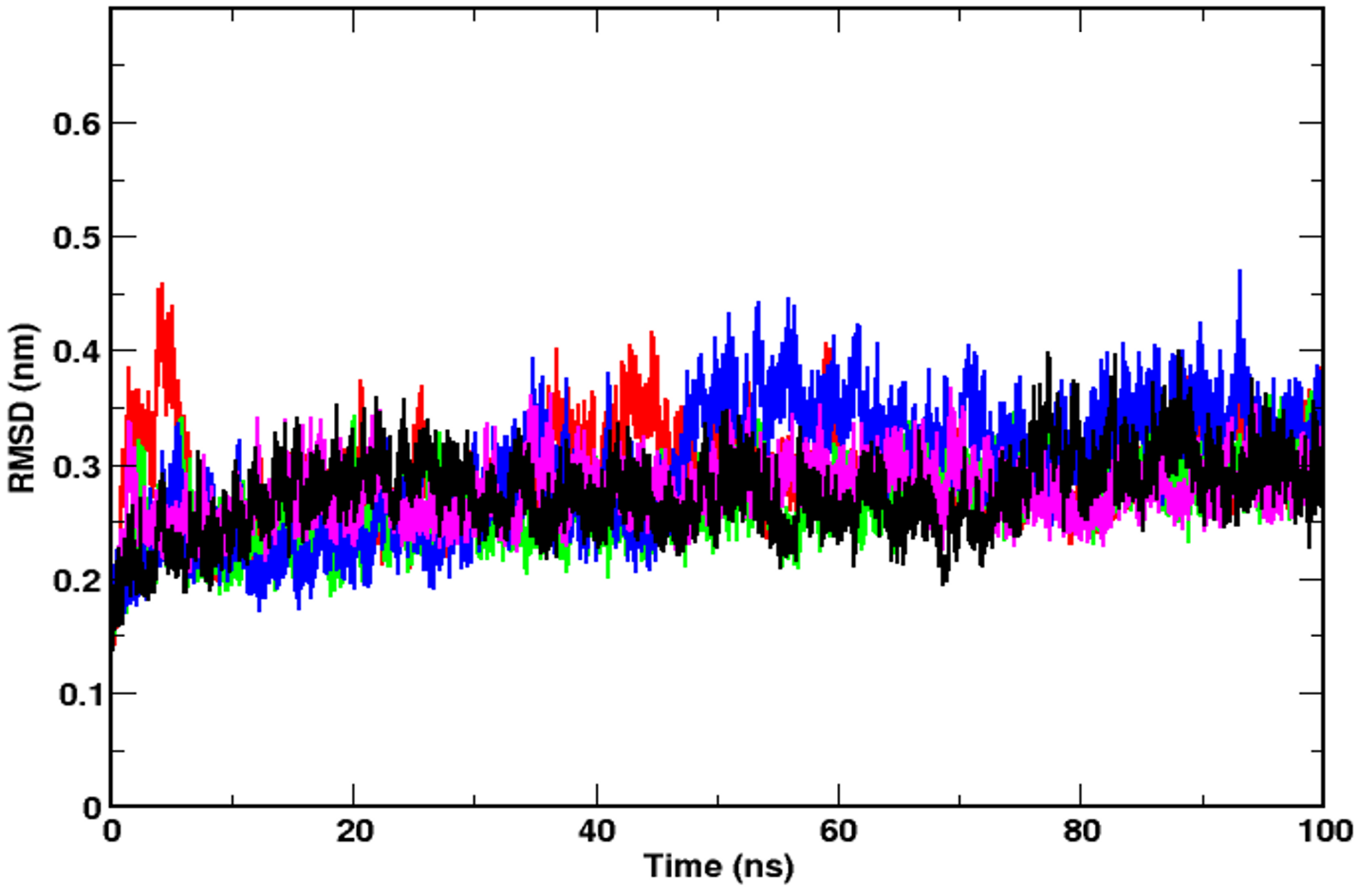

**Figure S3.** Time evolution of the RMSDs of the RBD/ACE2 protein complex in WT (black), Alpha (green), Beta (blue), Gamma (red) and Delta (magenta). The systems were found to be largely stable after 50ns.

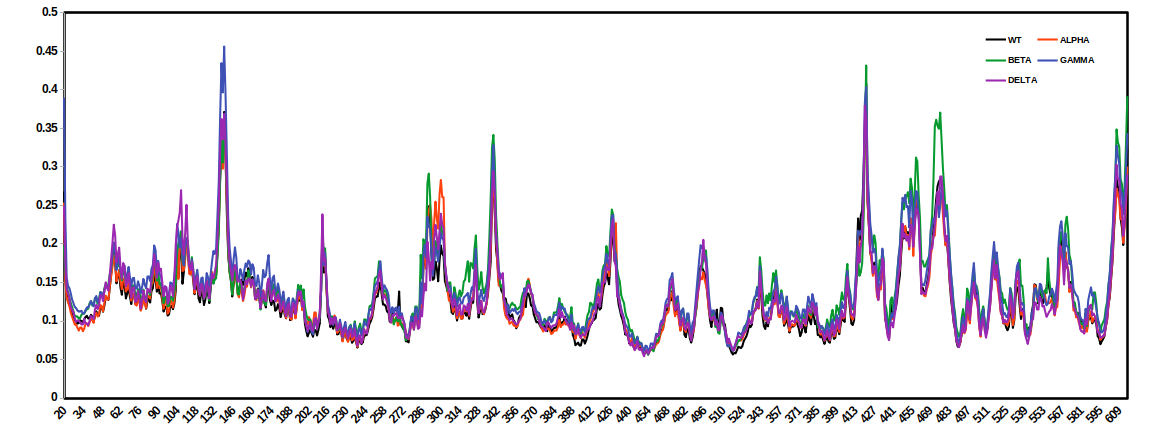

**Figure S4.** RMSF of CA atoms of all the systems under study.

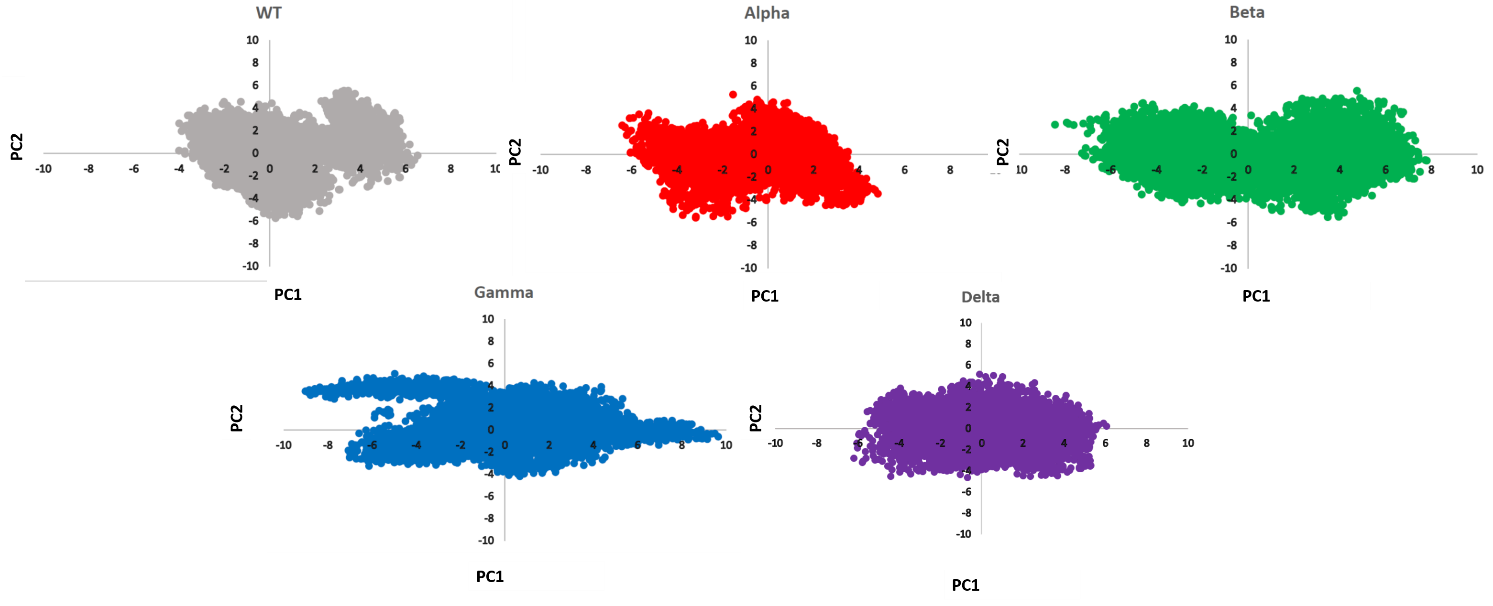

**Figure S5.** Distribution of simulated structures on the plane constituted by the first two principal components of ACE2 and RBD in WT (grey), Alpha (red), Beta (green), Gamma (blue) and Delta (magenta).

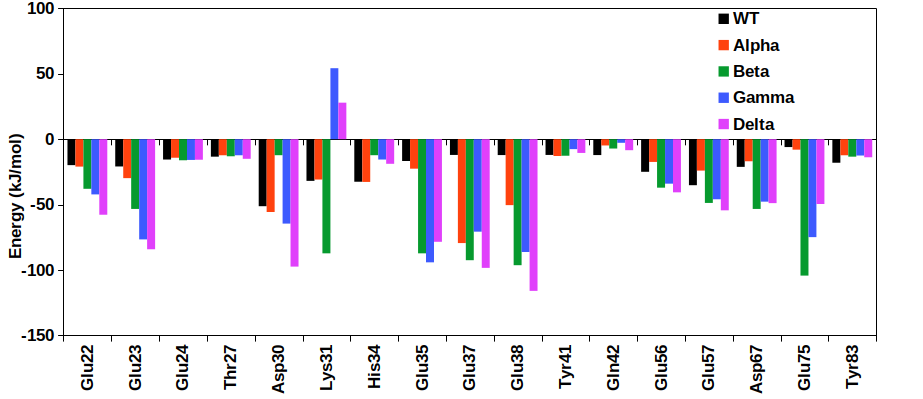

**Figure S6.** Residue level contribution of ACE2 towards the total interaction energy for all the five systems under study.

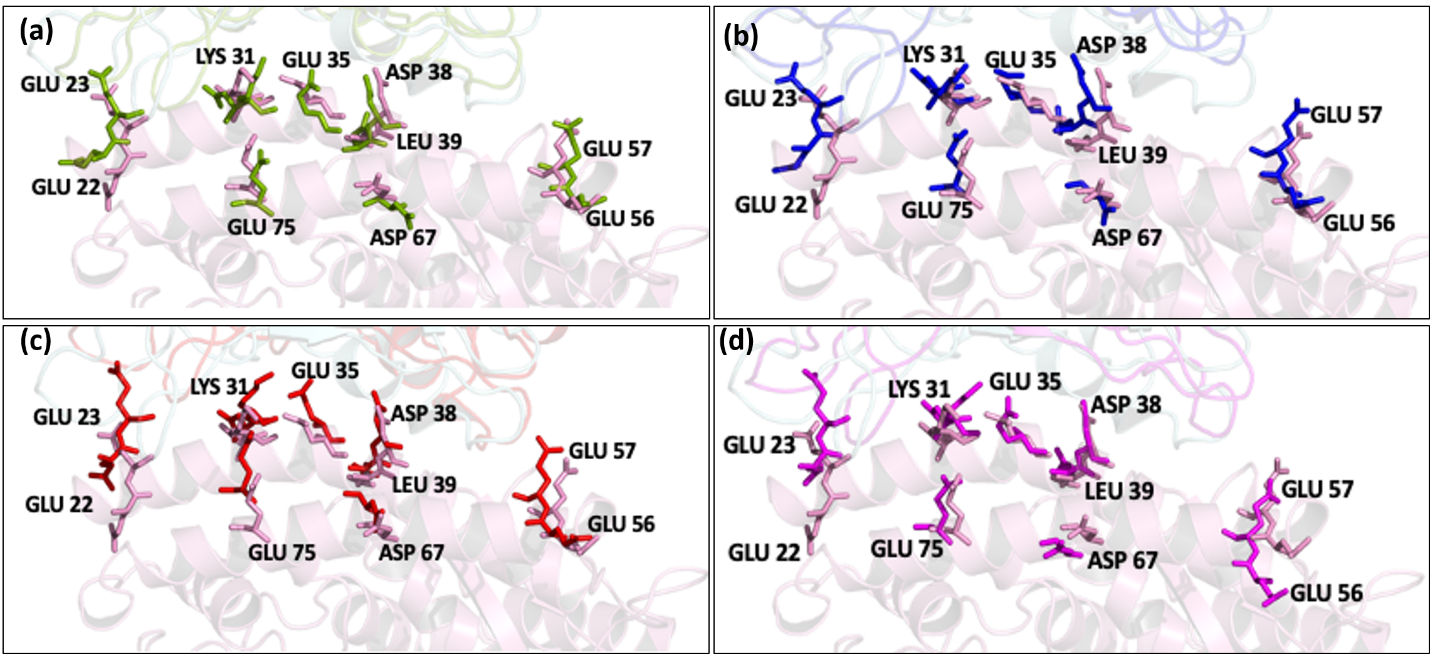

**Figure S7.** ACE2 interfacial residues that interact with RBD. Superimposed images of WT with (a) Alpha, (b) Beta (c) Gamma and (d) Delta. Color scheme: WT (pink), Alpha (green), Beta (blue), Gamma (red) and Delta (magenta). The residues lie at similar position in the ACE2 receptor.

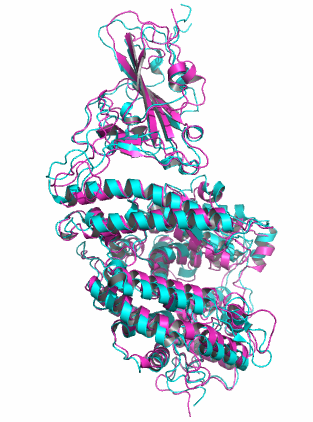

**Figure S8.** Superimposed structure of RBD/ACE2 complex of time-averaged B.167.2 structure (shown in magenta) and cryo-EM (PDB ID:78VB) (shown in cyan) [21] showing similarity between the two with RMSD 1.7 Å.

**Table S1. List of Systems**

| **Serial No** | **System** | **Simulation Time** |
| --- | --- | --- |
| 1 | WT | 100ns |
| 2 | Alpha (B.1.1.7 ) | 100ns |
| 3 | Beta (B.1.351) | 100ns |
| 4 | Gamma (P.1) | 100ns |
| 5 | Delta (B.1.167.2) | 100ns |

**Table S2. RMSD of the triplicates**

| **System** | **Average RMSD from triplicates** | **Std Deviation** |
| --- | --- | --- |
| WT | 0.273 | 0.03 |
| Alpha | 0.2718462 | 0.03481527 |
| Beta | 0.2707158 | 0.04382165 |
| Gamma | 0.2688205 | 0.04132388 |
| Delta | 0.2580426 | 0.02858201 |
